# Nuclear factor kappa B subunits and IkappaB family are modulated in the ovine endometrium during early pregnancy

**DOI:** 10.1101/2025.11.22.689904

**Authors:** Ziwang Du, Yanshu Xu, Haoran Yang, Haibei Zhu, Leying Zhang, Ling Yang

## Abstract

Pregnancy modulates the endometrial immune responses to establish maternal immune tolerance, and nuclear factor kappa B (NF-κB) subunits and IκB family are involved in the maternal innate and adaptive immune responses. It, nevertheless, is unclear if early pregnancy regulates the expression of NF-κB subunits and inhibitor of NF-κB (IκB) family in the ovine endometrium. In this study, ovine endometria were sampled at day 16 of the estrous cycle (N16), and at days 13, 16, and 25 of pregnancy (P13, P16, and P25), and mRNA and protein expression of NF-κB subunits and IκB family was analyzed by RT-qPCR, western blot, and immunohistochemistry. The results revealed that the expression of all NF-κB subunits was decreased at P25 compared with N16 both in mRNA and protein expression, but the expression of B cell leukemia-3, IκBα, IκB kinase γ, and IκBδ was increased at P25 compared with N16 both in mRNA and protein expression. IκBβ, however, was downregulated during early pregnancy, and the expression of IκBε and IκBζ was decreased at P13 and P16 compared with N16 and P25 both in mRNA and protein expression. In summary, early pregnancy changed the expression of NF-κB subunits and IκB family in the ovine endometrium both in mRNA and protein expression, which may be essential for maternal immune tolerance and embryo implantation.

## 1. Introduction

The endometrial receptivity plays a key role in embryo implantation, and uterine immune regulatory genes and immune response are essential for mediating uterine receptivity in humans [1]. Endometrial immune dysregulations, however, lead to the displaced window of implantation, which is related to endometrial immune profile and implantation failure in humans [2]. In addition, the reciprocating dialogue between the conceptus and cow is important for successful implantation, and immunological reactions at the uterus are involved in these processes during early pregnancy [3]. Furthermore, long non-coding RNAs and interferon-τ (IFNT)-responsive gene (interferon-stimulated gene 15-kDa protein, ISG15) expressed in the endometria are necessary for establishing endometrial receptivity and successful pregnancy in ruminants [4]. IFNT inhibits corpus luteum degeneration, allowing it to continuously secrete progesterone, and the synergistic interactions between progesterone and IFNT play a key role in establishing endometrial receptivity during pregnancy in ruminants [5]. Moreover, our previous reports indicate that the expression of nucleotide-binding domain-like receptors and toll-like receptors in the ovine endometria are modulated during early pregnancy, which are involved in the immunological dialogue between the uterus and conceptus [6,7]. It, therefore, is essential for exploring immune-related signal pathway in the endometria during early pregnancy.

Nuclear factor kappa B (NF-κB) transcription factors consist of NF-κB1 (*NFKB1*), NF-κB2 (*NFKB2*), RelA (*RELA*), RelB (*RELB*), and c-Rel (*REL*), which play an essential role in immunity and inflammation in mammals. Five NF-κB members have a conserved Rel homology domain in the N-terminal region, responsible for dimerization, binding to specific DNA regions and nuclear translocation [8]. In addition, NF-κB signaling is identified in pregnancy tissues, including myometrium, amnion, and cervical epithelium, which is crucial in human implantation, preterm labor prevention, and development of maternal tolerance to the fetus [9]. NF-κB signaling, however, is overactive in endometriotic lesions, which is involved in endometriosis development and recurrence in humans and mice [10]. Furthermore, as an initial pregnancy signal in ruminants, IFNT can inhibit endometrial inflammatory damage induced by lipopolysaccharide (LPS) through attenuating NF-κB signaling in bovine endometrial epithelial cells [11]. Moreover, IFNT treatment modulates NF-κB signaling in ovine uterine luminal epithelial cells, suggesting that NF-κB signaling is implicated in the physiological processes of pregnancy establishment [12].

NF-κB transcription factors interact with inhibitors of NF-κB (IκB) proteins, including IκBα (*NFKBIA*), IκBβ (*NFKBIB*), IκBε (*NFKBIE*), IκBδ (*NFKBIZ*), IκBδ (*NFKBID*), and B cell leukemia-3 (BCL-3), as well as NF-κB essential modulator (IκB kinase γ (IKKγ), also called *IKBKG*), and play a central role in coordinating immune and inflammation responses. In addition, IκB proteins in the cytoplasm can sequester NF-κB dimers to modulate the release of inflammatory cytokines and stress signals [8]. Furthermore, IκB family is expressed in the maternal spleen and lymph nodes, which is modulated by early pregnancy to participate in the establishment of maternal immune tolerance in ewes [13]. Downregulation of IκBα in endometrium, however, leads to the over-activation of NF-κB signaling, which increases the inflammatory state during decidualization and has a negative effect on embryo implantation in humans [14]. Moreover, a herbal formulation supplement improves endometrial receptivity and blastocyst implantation by inducing IκBα expression at uterus implantation sites in mice [15].

Conceptus signal (IFNT) and progesterone modulate the expression of implantation-related genes in the endometrium, which contributes to pregnancy establishment, conceptus survival, and elongation in ruminants [16]. In addition, the innate immune system in the uterus is modulated during early pregnancy, which is implicated in avoiding semi-allogeneic fetus rejected by the mother [5]. It, therefore, was hypothesized that NF-κB subunits and IκB family were regulated in the ovine endometria during early pregnancy. The objective of this study was to investigate the expression of mRNA and protein of NF-κB subunits and IκB proteins in the endometria from nonpregnant and early pregnant ewes, which will be helpful for understanding the establishment of maternal immune tolerance during early pregnancy.

## 2. Materials and methods

### 2.1. Animals and experimental design

All experimental procedures were approved by the Hebei University of Engineering Animal Care and Use Committee (HUEAE 2019-017). A total of 24 ewes (Small-tail Han sheep) with approximately 18 months of age, normal estrous cycles and similar body conditions (average weight of 41 kg, body condition score of 3) were received the same diet to meet the NRC (National Research Council, 2007) requirements, and used in the study. The females were housed under a condition of a 11 h light/13 h dark cycle, temperature of 3-18°C and free access to food and water. The animals were randomly divided into four different groups (n = 6 for each group) before breeding, including days 13 (P13), 16 (P16), and 25 (P25) of pregnancy, and day 16 of the estrous cycle (N16), as previously described [13]. There are significantly greater concentrations of progesterone in plasma on days 12-13, and lower concentrations of progesterone on days 15-16 during the ovine estrous cycle [17]. IFNT (Protein X) and additional proteins secreted by the trophoblast of blastocyst in the uterus are detected between days 14 and 21 [18], and IFNT reaches the maximum abundance on day 16 of pregnancy in sheep [19]. The specific stages, therefore, were selected as described above. A controlled internal drug release device was used for estrus synchronization. Estrus was detected by using vasectomized males, and ewes were mated with fertile rams (n = 3 for each group) in groups P13, P16, and P25. The ewes in group N16 were mated with vasectomized males. All ewes were slaughtered based on the pregnancy day or the estrous cycle designed in a local slaughterhouse, and the endometrial samples were taken from the ipsilateral endometrial horn to the corpus luteum. The tissue samples from all ewes were fixed in the 4% paraformaldehyde for 24 hours at room temperature, and also frozen in liquid nitrogen and stored at -80 ^°^C until further analysis (Figure 1).

**Figure 1.**
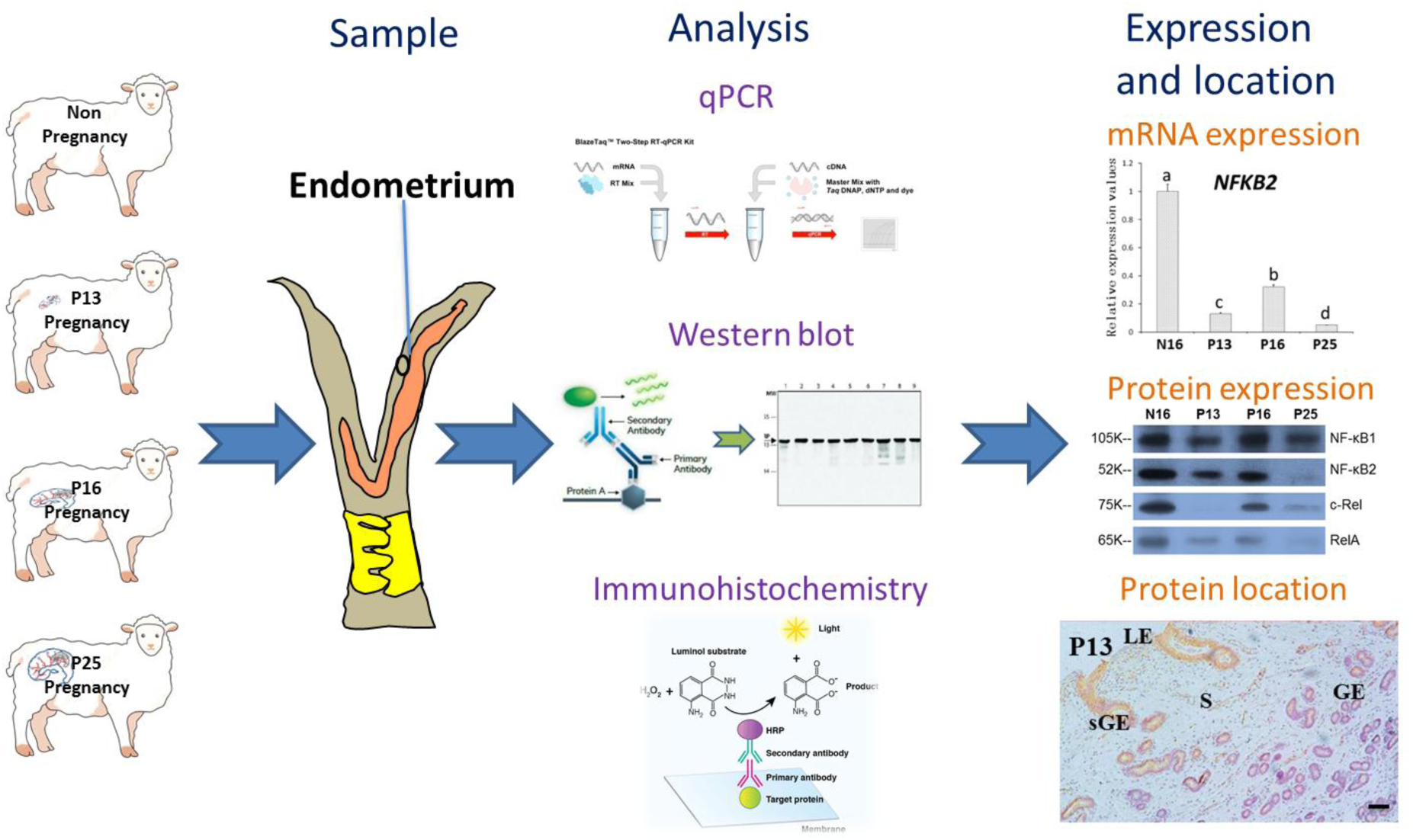
Overview of the experimental approach.

### 2.2. RNA extraction and RT-qPCR assay

The samples (transverse pieces) were crushed in liquid nitrogen, and total RNA was isolated using TRNzol Universal Reagent (DP424; Tiangen Biotech Co., Ltd., Beijing) according to the manufacturer’s instructions. The quantity and quality of total RNA were verified using agarose gel (1%) electrophoresis and spectrometry (A260/A280, NanoDrop 2000, Thermo Fisher Scientific, Waltham, MA, USA). The 260/280 ratios obtained from all samples were in the range of 1.9-2.1. A control that contained all the reaction components except for the reverse transcriptase was used to test for contaminating DNA. After the possible genomic DNA removal with gDNase, 1 μg of total RNA was used for cDNA synthesis according to manufacturer’s recommendations (FastQuant RT kit with gDNase, Tiangen Biotech). Quantity and quality of the cDNA were checked using the spectrometry and agarose gel electrophoresis. The specified primers listed in Table 1 were designed based on the sequences in the NCBI database (http://www.ncbi.nlm.nih.gov/) for the ovine genes of NF-κB subunits and the IκB family, and were synthesized by Shanghai Sangon Biotech Co., Ltd. Primer matrix experiments were performed with a range of concentrations from 50 nM to 900 nM in a logical series to determine the optimal primer concentrations, allowing the amplification efficiencies of the primer sequences in an acceptable range (between 0.9 and 1.1). The SuperReal PreMix Plus kit (Tiangen Biotech) was used for quantitative real-time PCR in a total volume of 20 μl (qPCR MasterMix, 10 μl; Forward primer (10 μΜ), 0.4 μl; Reverse primer (10 μΜ), 0.4 μl; cDNA, 1 μl; RNase-Free ddH_2_O, 8.2 μl) in triplicate using only H_2_O as no template control. The primer product was sequenced for checking specificity. PCR conditions were 40 cycles of 95 ^°^C for 10 sec, 60-62.5 ^°^C (60 ^°^C for *NFKB1, NFKB2,* and *BCL3*, 60.5 ^°^C for *NFKBIA*, *NFKBID* and *NFKBIZ*, 61 ^°^C for *REL*, *NFKBIB*, and *NFKBIE*, 62 ^°^C for *RELA*, *RELB*, and *IKBKG*) for 20 sec, and 72 ^°^C for 25 sec. Melting curve analysis, and electrophoresis on 2% agarose gel were used to verify PCR results. Glyceraldehyde phosphate dehydrogenase gene *(GAPDH*) was analyzed in parallel in all target genes, and used for normalization of gene expression data. Expression of *GAPDH* mRNA was checked and not affected by stage of gestation and pregnancy status. The 2^-ΔΔCt^ method [20] was employed to analyse relative expression value according to their cycle threshold using *GAPDH* as an internal control and normalizing by the samples from N16.

### 2.3. Western blot

The proteins of endometrial samples were extracted using a radio-immunoprecipitation assay (RIPA) lysis buffer with PMSF (Phenylmethylsulfonyl fluoride) (Biosharp, BL504A, Hefei, China), and the Bradford assay was used to measure the concentrations. The total proteins (10 μg/well) were separated on 12% sodium dodecyl sulfate-polyacrylamide gel electrophoresis, and then transferred electrophoretically to a 0.22 μm polyvinylidene fluoride membrane. The membranes were blocked with 5% fat-free milk at 4 ^°^C overnight, and then incubated with primary antibodies individually at 4 ^°^C overnight. The antibodies of NF-κB subunits and IκB proteins were provided in Table 2, and diluted with Tris-buffered saline with tween (TBST). The primary antibodies were validated through peptide blocking experiments and considered to be species cross-reactive by specific binding to native ovine proteins. After being washed with TBST, the membranes were incubated with secondary antibodies including goat anti-mouse IgG-HRP (Biosharp, BL001A, Hefei, China) or goat anti-rabbit IgG-HRP (Biosharp, BL003A) at a dilution of 1:2000 for one hour at room temperature, to detect the target proteins. An ECL (Enhanced Chemiluminescence) detection reagent (Tiangen Biotech) with X-ray films was used to reveal the protein bands. Band intensities were quantified using Quantity One V452 (Bio-Rad Laboratories) with a loading control by GAPDH antibody (Santa Cruz Biotechnology, sc-47724) in the same protocol.

### 2.4. Immunohistochemistry analysis

Paraformaldehyde-fixed tissues were embedded with paraffin, and cut into cross-sections of 5 μm. Some sections were stained with hematoxylin-eosin and examined under a light microscope (Nikon Eclipse E800, Japan). In addition, after antigen retrieval with boiling 0.01 M citric acid buffer (pH = 6.0), blocking endogenous peroxidase activity using 3% hydrogen peroxide and endogenous non-specific binding sites using 5% normal goat serum, other sections were incubated with primary antibodies, including RelA antibody (sc-8008, Santa Cruz Biotechnology, Santa Cruz, CA, USA) and c-Rel antibody and (sc-6955, Santa Cruz Biotechnology) for NF-κB subunits, as well as IκBα antibody (sc-1643, Santa Cruz Biotechnology) and IκBβ antibody (sc-390622, Santa Cruz Biotechnology) for IκB family, for one hour at room temperature at a 1:100 dilution with phosphate buffer saline (PBS). Negative controls were treated with host-specific IgG instead of the primary antibody at the same concentration. After being washed with PBS, a secondary antibody (goat anti-mouse IgG-HRP (Biosharp, BL001A) or goat anti-rabbit IgG-HRP (Biosharp, BL003A), at a dilution of 1:500 for one hour at room temperature, was used to detect the primary antibodies. A DAB kit (Tiangen Biotech) was used as the chromogen to detect the target proteins. Hematoxylin was used for counterstaining the tissue section. Sections were dehydrated with a series of increasing alcohol concentrations and sealed with a coverslip using neutral gum. Sections were examined with a light microscope (Nikon Eclipse E800, Japan), and images were captured and quantitative analysed by two investigators in a blinded fashion with covering labels, according to the following scale: 0, no staining; 1, weak staining; 2, strong staining; 3, stronger staining, as described previously [6].

### 2.5. Statistical analyses

The relative expression values of mRNA and protein for NF-κB subunits and IκB family were analyzed as a completely randomized design with six animals per group and four repeats for each ewe using the Proc Mixed models of SAS (Version 9.1; SAS Institute, Cary, NC). For endometria from different stage of gestation or pregnancy status, the model contained random effect of ewe and fixed effects of stage of gestation, pregnancy status and the interaction of stage of gestation and pregnancy status. Data normality was tested using the PROC UNIVARIATE procedure in SAS version 9.2 (SAS Institute Inc.), and the data were in normal distribution. The comparisons among the relative expression levels of different groups were made by using the Duncan’s multiple range post hoc test and controlling the experiment-wise type ± error equal to 0.05. Data are present as mean ± S.E.M. Statistical significance was defined as *P* < 0.05.

## 3. Results

### 3.1. Gene and protein expression of NF-κB subunits in the endometria

Figure 2A showed that there were declines in transcript abundances of *NFKB2*, *RELA*, and *REL* (*P* < 0.05) during early pregnancy (P13, P16, and P25) compared to N16, but expression values of *NFKB2* and *REL* were greater in P16 compared to P13 and P25 (*P* < 0.05). In addition, there was a downregulation of *RELA* in P25 compared to the other three groups (*P* < 0.05). Furthermore, *RELB* expression was downregulated in P16 and P25 compared to N16 and P13 (*P* < 0.05). The expression abundances of *NFKB1*, however, were greater in N16 and P16 compared to P13 and P25 (*P* < 0.05).

**Figure 2.**
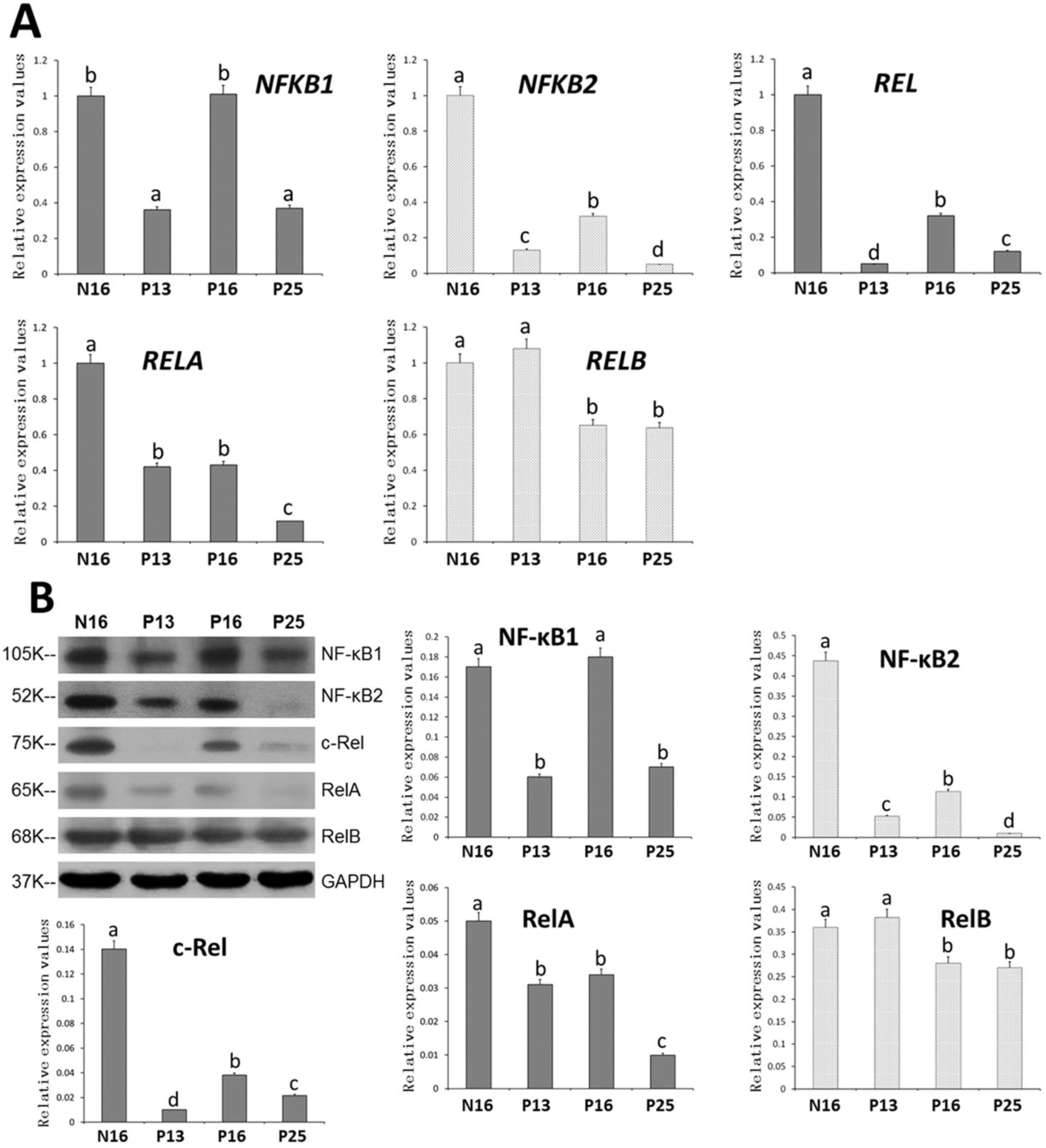
Expression of NF-κB subunits in the endometrium (n = 6 for each group). A: NF-κB subunit mRNA expression values. B: NF-κB subunit protein expression values. N16 = Day 16 of nonpregnancy; P13 = Day 13 of pregnancy; P16 = Day 16 of pregnancy; P25 = Day 25 of pregnancy; Significant differences (*P <* 0.05) are indicated by different letters.

Figure 2B indicated that the values of NF-κB1, NF-κB2, RelA, RelB, and c-Rel proteins in P25 were lower compared to N16 (*P* < 0.05). Early pregnancy inhibited the expression of NF-κB2, RelA, and c-Rel proteins, but NF-κB1, NF-κB2 and c-Rel protein expression was the greatest in P16 during early pregnancy (*P* < 0.05). NF-κB2 and RelA proteins were undetected in P25, and c-Rel protein was undetected in P13 and P25. RelA expression was greater in P13 and P16 compared to P25 (*P* < 0.05), and RelB protein expression was decreased in P16 and P25 compared to N16 and P13 (*P* < 0.05).

### 3.2. Immunohistochemistry for representative proteins of RelA and c-Rel in the endometria

Figure 3 indicated that RelA protein was strongly located in the uterine luminal epithelium (LE) and superficial glandular epithelium (sGE) with weakly staining in the glandular epithelium (GE) for groups N16, P13, and P16, and without detected for group P25. On the other hand, staining for c-Rel protein was mainly limited to superficial stroma and sGE for groups N16, P16, and P25, without detected for group P13. In addition, the staining intensities of the negative control, N16, P13, P16, and P25 were 0, 2, 1, 1, and 0 for RelA protein, as well as 0, 3, 0, 2, and 1 for c-Rel protein.

**Figure 3.**
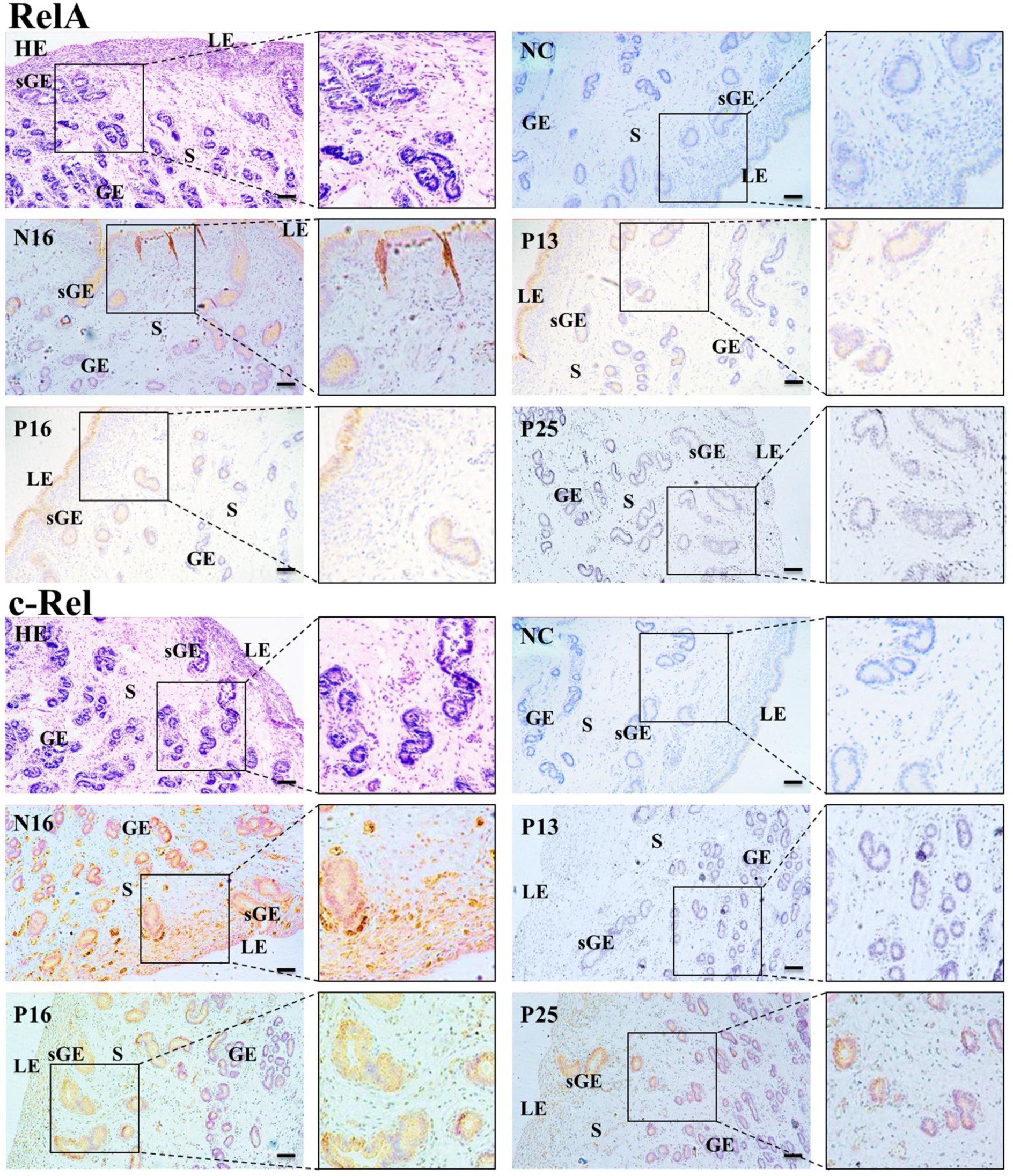
Representative immunohistochemical localization of the RELA and C-REL proteins in the endometrium (n = 6 for each group). NC = negative control; sGE = superficial glandular epithelium; GE = glandular epithelium; S = stroma; N16 = day 16 of the estrous cycle; P13 = day 13 of pregnancy; P16 = day 16 of pregnancy; P25 = day 25 of pregnancy. Bar = 100 µm.

### 3.3. Gene and protein expression of IκB family in the endometria

In the endometria, the expression abundances of *BCL3* and *NFKBID* mRNA were greater on P13, P16, and P25 compared to N16 (*P* < 0.05; Figure 4A). In addition, there was a peak in the expression values of *NFKBIA* and *IKBKG* in P25 compared to the other three groups (*P* < 0.05), but there was no significant difference among N16, P13, and P16 (*P* > 0.05). Early pregnancy, however, downregulated the expression of *NFKBIB*, with the lowest abundance in P16 (*P* < 0.05). Furthermore, transcript abundances of *NFKBIE* and *NFKBIZ* were lower in P13 and P16 compared to N16 and P25 (*P* < 0.05).

**Figure 4.**
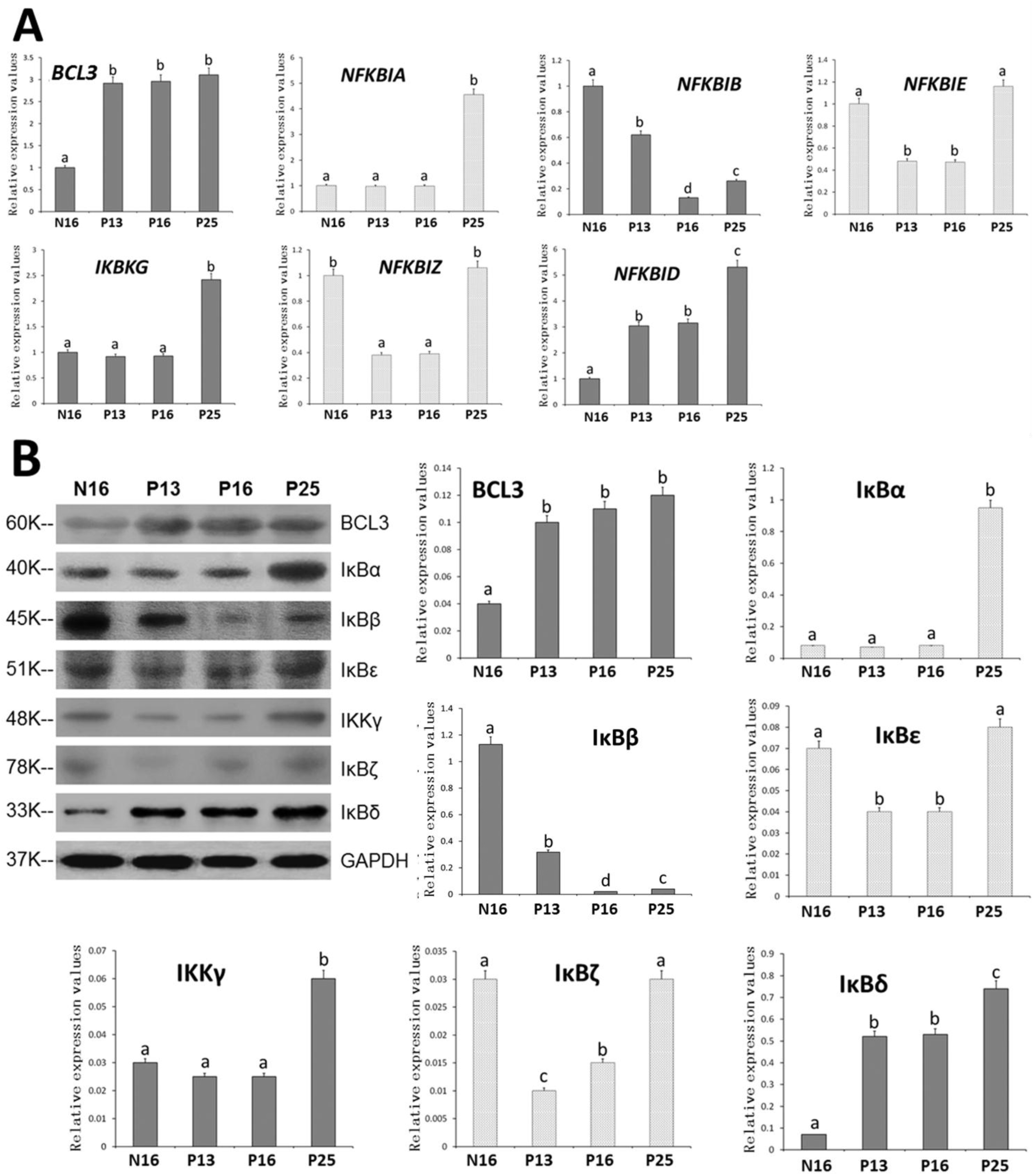
IκB family in the endometrium (n = 6 for each group). A: IκB family mRNA expression values. B: IκB family protein expression values. N16 = Day 16 of nonpregnancy; P13 = Day 13 of pregnancy; P16 = Day 16 of pregnancy; P25 = Day 25 of pregnancy; Significant differences (*P <* 0.05) are indicated by different letters.

Figure 4B showed that the protein expression of BCL-3, IκBα, IKKγ, and IκBδ was greater in P25 compared to N16 (*P* < 0.05), but early pregnancy suppressed the expression of IκBβ protein (*P* < 0.05). In addition, the protein expression of BCL3 and IκBδ was greater P13, P16, and P25 compared to N16 (*P* < 0.05), but protein expression of IκBε and IκBδ was lower in P13 and P16 compared to N16 and P25 (*P* < 0.05). There, however, was no significant difference among N16, P13, and P16 in IκBα and IKKγ proteins (*P* > 0.05), and IκBδ protein was undetected in P13.

### 3.4. Immunohistochemistry for representative proteins of IκBα and IκBβ in the endometria

It was revealed in Figure 5 that IκBα proteins were strongly located in the uterine sGE and with weakly staining in GE for groups N16, P13, P16, and P25. In addition, IκBβ protein was strongly expressed in the uterine sGE and superficial stroma with weakly staining in GE for groups N16, P13, and P25. Furthermore, the staining intensities of the negative control, N16, P13, P16, and P25 were 0, 1, 1, 1, and 2 for IκBα protein and 0, 3, 2, 0, and 1 for IκBβ protein.

**Figure 5.**
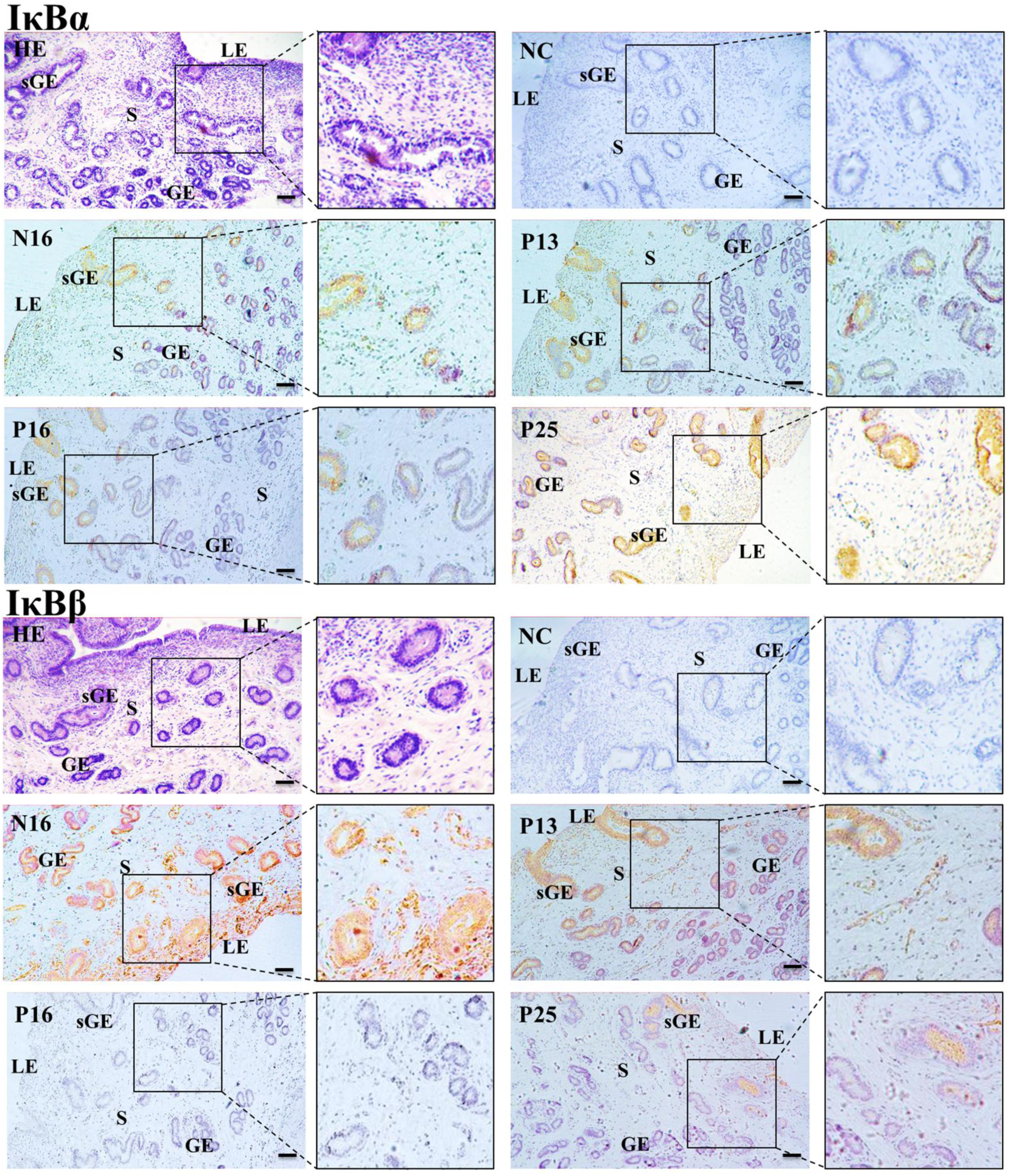
Representative immunohistochemical localization of the IκBα and IκBβ proteins in the endometrium (n = 6 for each group). NC = negative control; sGE = superficial glandular epithelium; GE = glandular epithelium; S = stroma; N16 = day 16 of the estrous cycle; P13 = day 13 of pregnancy; P16 = day 16 of pregnancy; P25 = day 25 of pregnancy. Bar = 100 µm.

## 4. Discussion

The endometrial immune responses are related to embryo implantation in humans [1], and the immunological reactions at the uterus play a key role in bovine successful pregnancy [3]. In addition, during early pregnancy in ewes, nucleotide-binding domain-like receptors and toll-like receptors are implicated in endometrial immune responses [6,7]. Furthermore, IFNT (early pregnancy recognition signals in ruminants) can modulate NF-κB signaling in bovine endometrial epithelial cells [11], and IκB proteins participate in embryo implantation in humans [14]. It, therefore, is suggested that early pregnancy exerts their effects on the endometria to modulate the expression of NF-κB subunits and IκB proteins, which may contribute to maternal immune tolerance and embryo implantation in sheep (Figure 6).

**Figure 6.**
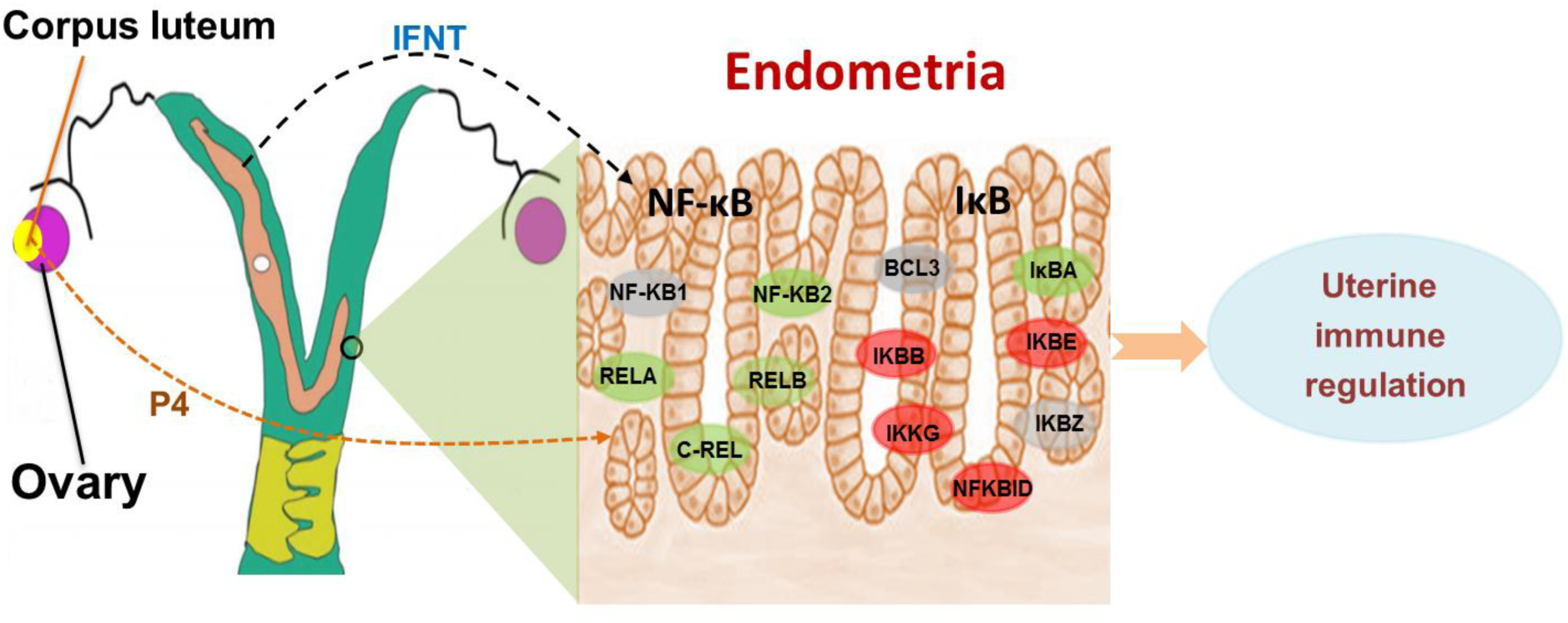
Sketch of NF-κB subunit and IκB family in the endometrium during early pregnancy. Early pregnancy induces changes in the expression of NF-κB subunits and IκB family, which may be involved in the maternal immune tolerance and embryo implantation. Red, stimulated; gray, changed; Green, inhibited.

In this study, early pregnancy inhibited mRNA and protein expression of endometrial NF-κB1 (*NFKB1*) and RelA (*RELA*) in the ovine endometria, and RelA protein was strongly located in the uterine LE and sGE. It has been reported that expression values of endometrial NF-κB1 and RelA are greater in the women with recurrent implantation failure compared with the normal control group [21]. In addition, the protein expression of NF-κB1 and RelA is greater in women with endometrial polyps than in fertile controls, which are associated with lower endometrial receptivity and sub-fertility [22]. Furthermore, early pregnancy suppresses the NF-κB1 expression in the maternal liver, which is involved in maternal hepatic homeostasis and immune tolerance in the ovine [23]. Moreover, IFNT can induce maternal immune tolerance, and attenuate LPS-induced endometrial inflammatory damage through inhibiting RelA expression in the bovine endometrial epithelial cells [11]. On the other hand, early pregnancy downregulates RelA expression in the maternal thymus, which is related to the establishment of maternal immune tolerance and successful pregnancy in ewes [24]. It, therefore, is suggested that the downregulation of endometrial NF-κB1 and RelA may be beneficial for the establishment of maternal immune tolerance during early pregnancy in ewes.

Our data indicated that there was a decline in mRNA and protein expression of NF-κB2 (*NFKB2*) and RelB (*RELB*) in the ovine endometria. It has been reported that a greater expression abundance of *NFKB2* mRNA transcript in pig uterine horns induced by is related to the uterine inflammatory response [25]. In addition, the upregulation of *NFKB2* mRNA in bovine uterine epithelial cells stimulated by great concentration of sperm is involved in sperm phagocytosis by polymorphonuclear neutrophils [26]. Furthermore, NF-κB2 and RelB are related to the corticotropin-releasing hormone produced in the full-term human placenta [27], and the upregulation of NF-κB2 and RelB in myeloid-derived suppressor cells induced by *Toxoplasma gondii* infection has a negative effect on maintaining maternal-fetal tolerance in mice [28]. It, therefore, suggested that the downregulation of endometrial NF-κB2 and RelB may be beneficial for the establishment of maternal-fetal tolerance during early pregnancy in ewes.

Our results showed that the c-Rel protein was mainly limited to superficial stroma and sGE, and c-Rel protein and mRNA expression was decreased in the endometria during early pregnancy with a slight increase at P16. It has been reported that the downregulation of c-Rel in the myometrium and myometrial biopsies plays a key role in myometrial quiescence and pregnancy maintenance in humans [29,30]. In addition, the downregulation of c-Rel in inguinal lymph nodes is related to maternal immune tolerance during early pregnancy in sheep [31]. It, therefore, is suggested that the downregulation of endometrial c-Rel during early pregnancy may contribute to decreasing endometrial inflammation, and the slight increase at P16 may be related to the peak of IFNT in ewes.

This study revealed that mRNA and protein expression of BCL3 and IκBδ (*NFKBID*) was upregulated during early pregnancy. It has been reported that BCL3 is an inhibitor of the NF-κB activity, and participates in the suppression of the innate immune response in mice [9]. In addition, the upregulation of BCL3 in the uterus of mice during diestrus is implicated in preventing inflammation [32], and the upregulation of *BCL3* mRNA and protein in the maternal spleen and thymus is involved in regulating splenic and thymic adaptive immunity and maternal immune tolerance in sheep [13,33]. Furthermore, the upregulation of IκBδ in the uterine tissue during implantation/placentation can restrict embryo arrestment in pregnant mice [34]. Moreover, the dynamic regulation of NF-κB activity by IκBδ in the uterus contributes to blastocyst implantation in mice [32]. The upregulation of endometrial BCL3 and IκBδ during early pregnancy, therefore, may contribute to embryo implantation and establishment of the maternal immune tolerance in ewes.

It was showed in this study that early pregnancy enhanced mRNA and protein expression of IκBα (*NFKBIA*) at P25 but suppressed mRNA and protein expression of IκBβ (*NFKBIB*), especially at P16 in the endometria. In addition, IκBα and IκBβ proteins were strongly located in the uterine sGE and with weakly staining in GE, and IκBβ protein was also strongly expressed in superficial stroma. It has been reported that IκBα can inhibit the activation of NF-κB to exert a beneficial effect on reducing the symptoms of preeclampsia in rats [35]. Furthermore, the degradation of IκBα in human peripheral blood mononuclear cells is associated with spontaneous preterm birth [36], but the upregulation of IκBα in the maternal spleen and liver is essential for the modulation of maternal splenic and hepatic functions and pregnancy establishment in sheep [13,37]. Moreover, the downregulation of IκBβ in peripheral blood mononuclear cells and endometrium plays a key role in the implantation process in humans [38,39]. It, therefore, is suggested that the upregulation of IκBα at P25 may be related to pregnancy establishment and maintenance. The downregulation of IκBβ, however, may be involved in the implantation process, and the deep decrease at P16 may be related to the peak of IFNT in ewes.

Protein and mRNA of endometrial IκBε (*NFKBIE*) and IκBδ (*NFKBIZ*) was downregulated at P13 and P16 in this study. It has been reported that *NFKBIE* (encoding IκBε) mutations in chronic lymphocytic leukemia result in immune escape in murine models [40], and IκBε is involved in glucose metabolism in endometrial cells [41]. In addition, IκBδ is involved in maternal immune tolerance in humans [42], and there is a negative relationship between IκBδ production and type I interferon secretion [43]. It, therefore, is suggested that the changes in expression of IκBε and IκBδ may be associated with endometrial immune modulation and glucose metabolism, and the downregulation of IκBδ at P16 may be related to the production of type I interferon (IFNT) from the conceptus.

The data of this study showed that IKKγ (*IKBKG*) expression of mRNA and protein was increased during early pregnancy. It has been reported that *IKBKG* gene expression (encoding IKKγ) is greater in the placentas of healthy controls compared with the women with preeclampsia, suggesting that IKKγ can inhibit the development of preeclampsia in humans [44]. In addition, *IKBKG* gene variants in the maternal genomes enhance the likelihood of preeclampsia [45], but a gradual increase in *IKBKG* mRNA and protein expression in the maternal spleen during early pregnancy is related to the immune regulation of the maternal spleen in ewes [13]. It, therefore, suggested that the increasing expression of the IKKγ in endometria during early pregnancy may be helpful for pregnancy establishment in ewes.

## 5. Conclusion

This study revealed for the first time that early pregnancy changed the expression of NF-κB subunits and IκB family in the ovine endometria. Compared with N16, the expression of all NF-κB subunits was decreased at P25, but the expression of BCL3, IκBα, IKKγ, and IκBδ was increased at P25 compared with N16, which may contribute to maternal immune tolerance. In addition, during early pregnancy, the expression of NF-κB1, NF-κB2, and c-Rel was upregulated, but IκBβ expression was downregulated at P16, which may be related to the peak of IFNT secretion. The downregulation of IκBβ during early pregnancy, and decreased expression of IκBε and IκBδ at P13 and P16, however, may be associated with the secretion of IFNT, progesterone, and other pregnancy signals, which participate in embryo implantation in ewes. Further researches are needed to confirm the mechanism of maternal immune tolerance and embryo implantation.

## CRediT authorship contribution statement

**Ziwang Du, Yanshu Xu** and **Haoran Yang**: Investigation. **Haibei Zhu**: Validation. **Leying Zhang**: Writing - original draft. **Ling Yang**: Conceptualization, Writing original draft, Writing – review & editing.

## Acknowledgments

This work was supported by the grants from the Natural Science Foundation of Hebei Province, China (C2024402023).

